# Constructing Functional Networks of Phosphorylation Sites Using Co-Phosphorylation

**DOI:** 10.1101/2021.08.16.456448

**Authors:** Marzieh Ayati, Serhan Yılmaz, Mark R. Chance, Mehmet Koyutürk

**Affiliations:** Deparment of Computer Science, University of Texas - Rio Grande Valley; Department of Computer and Data Sciences, Case Western Reserve University; Center for Proteomics and Bioinformatics, Case Western Reserve University; Department of Nutrition, Case Western Reserve University

## Abstract

Protein phosphorylation is a ubiquitous regulatory mechanism that plays a central role in cellular signaling. According to recent estimates, up to 70% of human proteins can be phosphorylated. Therefore, characterization of phosphorylation dynamics is critical for understanding a broad range of biological and biochemical processes. Technologies based on mass spectrometry are rapidly advancing to meet the needs for high-throughput screening of phosphorylation. These technologies enable untargeted quantification of thousands of phosphorylation sites in a given sample. Many labs are already utilizing these technologies to comprehensively characterize signaling landscapes by examining perturbations with drugs and knockdown approaches, or by assessing diverse phenotypes in cancers, neuro-degerenational diseases, infectious diseases, and normal development. Here, we comprehensively investigate the concept of “co-phosphorylation”, defined as the correlated phosphorylation of a pair of phosphosites across various biological states. We integrate nine publicly available phospho-proteomics datasets for various diseases (including breast cancer, ovarian cancer and Alzhemier’s disease) and utilize functional data related to sequence, evolutionary histories, kinase annotations, and pathway annotations to investigate the functional relevance of co-phosphorylation. Our results across a broad range of studies consistently show that functionally associated sites tend to exhibit significant positive or negative co-phosphorylation. Specifically, we show that co-phosphorylation can be used to predict with high precision the sites that are on the same pathway or that are targeted by the same kinase. Overall, these results establish co-phosphorylation as a useful resource for analyzing phospho-proteins in a network context, which can help extend our knowledge on cellular signaling and its dysregulation.

## 1 Introduction

Protein phosphorylation is a ubiquitous mechanism of post-translational modification observed across cell types and species. Recent estimates suggest that up to 70% of cellular proteins can be phosphorylated [1]. Phosphorylation is regulated by networks composed of kinases, phosphatases, and their substrates. Characterization of these networks is increasingly important in many biomedical applications, including identification of novel disease-specific drug targets, development of patient-specific therapeutics, and prediction of treatment outcomes [2, 3].

Phosphorylation is particularly important in the context of cancer, as down-regulation of tumor suppressors and up-regulation of oncogenes (often kinases themselves) by dysregulation of the associated kinase and phosphatase networks are shown to have key roles in tumor growth and progression [4, 5]. To this end, characterization of signaling networks enables exploration of the interconnected targets [6,7, 8] and identification of causal pathways [9], leading to the development of kinase inhibitors to treat a variety of cancers [10, 11]. Disruptions in the phosphorylation of various signaling proteins have also been implicated in the pathophysiology of various other diseases, including Alzheimer’s disease [12], Parkinson’s disease [13], obesity and diabetes [14], and fatty liver disease [15], among others. As a consequence, there is increased attention to cellular signaling in biomedical applications, motivating researchers to study phosphorylation at larger scales [16].

In response to the growing need for large-scale monitoring of phosphorylation, advanced mass spectrometry (MS)- based phospho-proteomics technologies have exploded [17]. These technologies enable simultaneous identification and quantification of thousands of phosphopeptides and phosphosites from a given sample [18]. These developments result in the generation of data representing the phosphorylation levels of hundreds of thousands of phosphosites under various conditions across a range of biological contexts, including samples from human patients, cell lines, xenografts, and mouse models [19]. As compared to the widespread availability and sharing of genomic and transcriptomic data, public repositories of phospho-proteomic data are sparse, but growing. As a consequence, secondary or integrative analyses of phospho-proteomic data are less common. Despite tremendous advances in the last decade, a majority of the human phosphoroteome has not been annotated to date [20]. Technical issues such as noise, lower coverage, lower number of samples, and low overlap between studies further complicate the analysis of phospho-proteomic data from a systems biology perspective [19].

In order to facilitate large-scale utilization of phospho-proteomic data, we introduced the notion of co-phosphorylation (Co-P) [21]. The motivation behind this approach is to represent phosphorylation data in the form of relationships between pairs of phosphosites. Defining co-phosphorylation as the correlation between pairs of phosphosites across a range of biological states within a given study, we alleviate such issues as batch effects between different studies and missing identifications, while integrating phosphorylation data across multiple studies. Recently, we applied Co-P to the prediction of kinase-substrate associations) [21] and unsupervised identification of breast cancer subtypes [22], showing that co-P enables effective integration of multiple datasets and enhances the reproducibility of predictions.

Co-phosphorylation is similar in spirit, but distinct and complementary to the notion of co-occurrence [23]. Co-occurence qualitatively assesses the relationship between the identification patterns of phosphosites in a broad range of studies. Co-P, on the other hand, quantitatively assesses the relationship between the phosphorylation levels of sites across a set of biological states (within a single study or by integrating different studies). Thus, co-occurrence captures high-level functional associations among phosphosites, whereas Co-P can also discover context-specific associations and provide insights into the dynamics of signaling interactions.

In this paper, we comprehensively characterize the relationship between co-phosphorylation and functional associations/interactions among protein phosphorylation sites. For this purpose, we systematically compare Co-P networks to networks that represent other functional relationships between proteins and phosphosites. These analyses serve two purposes: (i) Validation of Co-P as a relevant and useful tool for inferring functional relationships between proteins, (ii) Generation of knowledge on the basic principles of post-translational regulation of proteins and the functional relationships between them.

## 2 Materials and Methods

### 2.1 Phospho-Proteomic Datasets

We analyze 9 different MS-based phospho-proteomics data representing cancer and non-cancer diseases.

- **BC1 (Breast Cancer):** Huang et al. [24] used the isobaric tags for relative and absolute quantification (iTRAQ) to identify 56874 phosphosites in 24 breast cancer PDX models.
- **BC2 (Breast Cancer):** This dataset was generated to analyze the effect of delayed cold Ischemia on the stability of phosphoproteins in tumor samples using quantitative LC-MS/MS. The phosphorylation level of the tumor samples was measured across 3 time points [25]. The dataset includes 8150 phosphosites mapping to 3025 phosphoproteins in 18 breast cancer xenografts.
- **BC3 (Breast Cancer):** The NCI Clinical Proteomic Tumor Analysis Consortium (CPTAC) conducted an extensive MS based phosphoproteomics analysis of TCGA breast cancer samples [26]. After selecting the subset of samples to have the highest coverage and filtering the phosphosites with missing intensity values in those tumors, the remaining data contained intensity values for 11018 phosphosites mapping to 8304 phosphoproteins in 20 tumor samples.
- **OC1 (Ovarian Cancer):** This dataset was generated by the same study as BC2, using the same protocol. The dataset includes 5017 phosphosites corresponding to 2425 phosphoproteins in 12 ovarian tumor samples.
- **OC2 (Ovarian Cancer):** The Clinical Proteomic Tumor Analysis Consortium conducted an extensive MS based phosphoproteomic of ovarian HGSC tumors characterized by The Cancer Genome Atlas [27]. We filtered out the phosphosites with missing data and also selected a subset of tumors to maximize the number of phosphosites. This resulted in a total of 5017 phosphosites from 2425 proteins in 12 tumor samples.
- **CRC (Colorectal Cancer):** Abe et al. [28] performed immobilized metal-ion affinity chromatography-based phosphoproteomics and highly sensitive pY proteomic analyses to obtain data from 4 different colorectal cancer cell line. The dataset included 5357 phosphosites with intensity values cross 12 different conditions. These phosphosites map to 2228 phosphoproteins.
- **LC (Lung Cancer):** Wiredja et al. [29] performed a time course label-free phospho-proteomics on non-small lung cancer cell lines across 1, 6 and 24 hrs after applying two different treatments of PP2A activator and MK-AZD, resulting in total of 6 samples. They reported phosphorylation levels for 5068 phosphosites, which map to 2168 proteins.
- **AD (Alzheimer’s Disease):** LC-MS/MS phosphoproteomics was performed on eight individual AD and eight age-matched control postmortem human brain tissues. The dataset contains 5569 phosphosites mapping to 2106 proteins [30].
- **RPE (Retinal Pigmented Epithelium):** MS-based phosphoproteomics was performed on three cultured human-derived RPE-like ARPE-19 cells which were exposed to photoreceptor outer segments (POS) for different time periods (0, 15, 30, 60, 90, and 120 min) [31]. The dataset contains 1016 phosphosites mapping to 619 proteins in 18 samples.

### 2.2 Functional Networks

To assess the functional relevance of co-phosphorylation, we use networks of functional relationships/associations between pairs of phosphorylation sites. For this purpose, we consider four types of functional networks:

#### Kinase-Substrate Associations (KSAs)

We use PhosphoSitePLUS (PSP) [32] as a gold-standard dataset for kinase-substrate associations. PSP reports 9699 associations among 347 kinases and 6906 substrates. We use these associations to constructed a “shared kinase network” of phosphorylation sites, in which nodes represent phosphosites and edges represent the presence of at least one kinase that phosphorylated both sites. The associations obtained from PSP lead to a shared kinase network of 6906 phosphosite nodes and 881685 shared kinase edges.

#### Protein-Protein Interaction (PPI)

We use the PPIs that are provided in STRING database [33] with high confidence (combined score≥0.95). Overall, there are 61452 high-confidence interactions among 8987 proteins. For each of the 9 datasets, we use these PPIs to construct an interaction network among the sites identified in that dataset. In this network, each node represents a phosphosite and each edge represents an interaction between the two proteins that harbor the respective sites.

#### Evolutionary and Functional Associations

PTMCode is a database of known and predicted functional associations between phosphorylation and other post-translational modification sites [34]. The associations included in PTMCode are curated from the literature, inferred from residue co-evolution, or are based on the structural distances between phosphosites. We utilize PTMcode as a direct source of functional, evolutionary, and structural associations between phosphorylation sites. In the PTMcode network, there are 96519 phosphosite nodes and 4661285 functional association edges between these phosphosites.

#### Phosphosite-Specific Signaling Pathways

We use PTMsigDB as a reference database of site-specific phosphorylation signatures of kinases, perturbations, and signaling pathways [35]. While PTMSigDB provides data on all post-translational modifications, we here use the subset that corresponds to phosphorylation. There area 2398 phosphosites that are associated with 388 different perturbation and signaling pathways. We represent these associations as a binary network of signaling-pathway associations among phosphosites, in which an edge between two phosphosites indicates that the phosphorylation of the two sites is involved in the same pathway. The resulting network contains 6276 edges between 2398 phosphosite nodes.

For each functional network, the number of nodes/edges edges that overlap with our 9 phospho-proteomic datasets are shown in Table 1.

**Table 1:** Descriptive statistics of the phospho-proteomic datasets used in our computational experiments and their overlap with functional networks. For each dataset, the number of samples, the number of phosphorylation sites that were identified and the number of proteins that are spanned by these sites are shown. For each dataset and functional network pair, the number in the first row shows the number of sites with at least one interaction in the functional network and the second row shows the number of interactions in the functional network with both sites present in the corresponding dataset.

| Dataset | Descriptive Statistics |  |  | Overlap with Functional Networks |  |  |  |
| --- | --- | --- | --- | --- | --- | --- | --- |
|  | # Samples | # Phosphosites | # Proteins | Shared Kinase | PPI | PTMCode | PTMSigDB |
| BC1 | 24 | 15780 | 4539 | 805<br>27791 | 7632<br>142077 | 4437<br>15335 | 138<br>2547 |
| BC2 | 18 | 8150 | 3025 | 243<br>2723 | 1639<br>16541 | 1007<br>1811 | 54<br>429 |
| BC3 | 20 | 11472 | 3312 | 553<br>13123 | 4491<br>45911 | 3014<br>9127 | 119<br>2226 |
| OC1 | 12 | 5017 | 2425 | 414<br>7174 | 2450<br>17584 | 1318<br>2580 | 74<br>1032 |
| OC2 | 12 | 4802 | 2230 | 157<br>1114 | 818<br>4764 | 510<br>685 | 32<br>158 |
| CRC | 12 | 5352 | 2228 | 320<br>6237 | 1663<br>17573 | 1240<br>2715 | 51<br>421 |
| LC | 6 | 5068 | 2168 | 380<br>6493 | 2036<br>17884 | 1238<br>2919 | 64<br>588 |
| AD | 8 | 5569 | 1559 | 238<br>3637 | 1743<br>19075 | 941<br>3182 | 44<br>228 |
| RPE | 18 | 1016 | 619 | 120<br>931 | 371<br>1667 | 193<br>320 | 31<br>216 |

### 2.3 Assessment of Co-Phosphorylation

For a given phosho-proteomic dataset, we define the vector containing the phosphorylation levels of a phosphosite across a number of biological states as the *phosphorylation profile* of a phosphosite. For a pair of phosphosites, we define the co-phosphorylation of the two sites as the statistical association of their phosphorylation profiles. To assess statistical association, we refer to the rich literature on the assessment of gene co-expression based on mRNA-level gene expression [36], and consider Pearson correlation [37], biweight-midcorrelation [38], and mutual information [39]. Since our experiments suggest that the different measures of association lead to similar results (data not shown), we use Pearson correlation as a simple measure of statistical association in our experiments.

We use the datasets described in the previous section to characterize co-phosphorylation in relation to the functional, structural, and evolutionarily relationships between sites and proteins encoded in the functional networks. For this analysis, we investigate correspondence between co-phosphorylation in each individual MS-based phospho-proteomics dataset and each functional network.

### 2.4 Integration of Co-Phosphorylation Networks Across Datasets

Since co-phosphorylation can potentially capture context-specific, as well as universal functional relationships among phosphorylation sites, we also investigate the functional relevance of co-phosphorylation across different datasets. While integrating co-phosphorylation across multiple datasets, the number of samples (i.e., the number of dimensions used to compute the correlation) in each dataset is different. For this reason, we use the adjusted *R*-squared [40] (denoted 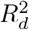) to remove the effect of number of dimensions from dataset-specific co-phosphorylation between pairs of phosphosites:

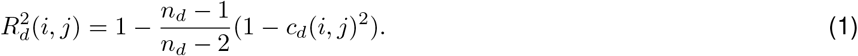

Here, *c_d_*(*i, j*) denotes the co-phosphorylation (measured by Pearson correlation) in dataset *d* ∈ *D* with *n_d_* samples.

In mass-spectrometry based phospho-proteomics, the overlap between the phosphorylation sites that are identified across different studies is usually low [19]. Specifically, for the 9 datasets we use in our computational experiments, thereare only17phosphosites thatare identified in all studies. Consequently, to preserve the scope of our cross-dataset analysis, we use all sites that are identified in at least one study. For this purpose, we develop a measure of cross-dataset co-phosphorylation that can integrate the co-phosphorylation scores computed on an arbitrary number of datasets. To handlemissingdatawithoutintroducingbias, we set 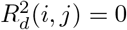 if phosphosite *i* or phosphosite *j* is not present in dataset *d*. Subsequently, we compute the integrated Co-P between sites *i* and *j* as follows:

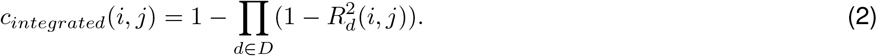

Observe that, 0 ≤ *c_integrated_*(*i, j*) ≤ 1, where the minimum value is realized if the two sites are never identified in the same dataset or their phosphorylation levels have zero correlation if they are identified together. If the phosphorylation levels of two sites exhibit perfect correlation in at least one dataset, then *c_integrated_* =1. Finally, as the number of datasets on which both sites are identified goes up, *c_integrated_* also tends to go up. Thus *c_integrated_* can be thought of as a measure of both co-occurrence [23]and co-phosphorylation [21], since it captures both the tendency of the sites being identified in similar contexts, as well as the relationship between their dynamic ranges of phosphorylation.

## 3 Results and Discussion

### 3.1 Statistical Significance of Co-phosphorylation

To understand whether the notion of co-phosphorylation (co-P) is biologically relevant, we first investigate the distribution of co-P levels across all pairs of phosphosites identified within a study. The results of this analysis for 9 datasets are shown in Figure 1. As seen in the figure, co-P follows a normal distribution with mean close to zero (as would be expected if phosphorylation levels were drawn from a normal distribution) and the distribution is narrower (and likely less noisy) if more biological states (dimensions) are available. Based on the premise that co-P can capture functionally relevant relationships, we hypothesize that distribution of co-phosphorylation on real datasets would contain more positively and negatively correlated phosphosite pairs than would be expected at random. To test this hypothesis, we conduct permutation tests by permuting phosphorylation levels across the entire data matrix, and compute the co-P distribution on these randomized datasets. As seen in the figure, co-P is concentrated more on strongly positive or strongly negative correlation levels for all datasets. For all datasets, the Kolmogorov-Smirnov (KS) test p-values for the difference between the observed co-P distribution and permuted co-P of distribution are ≪ 1*E* - 9. Similarly, the t-test *p*-values for the difference between the means of these distributions are ≪ 1*E*-9 for all datasets except CRC. The mean difference and the 95% confidence interval for each dataset are provided below the histograms in the figure.

**Figure 1:**
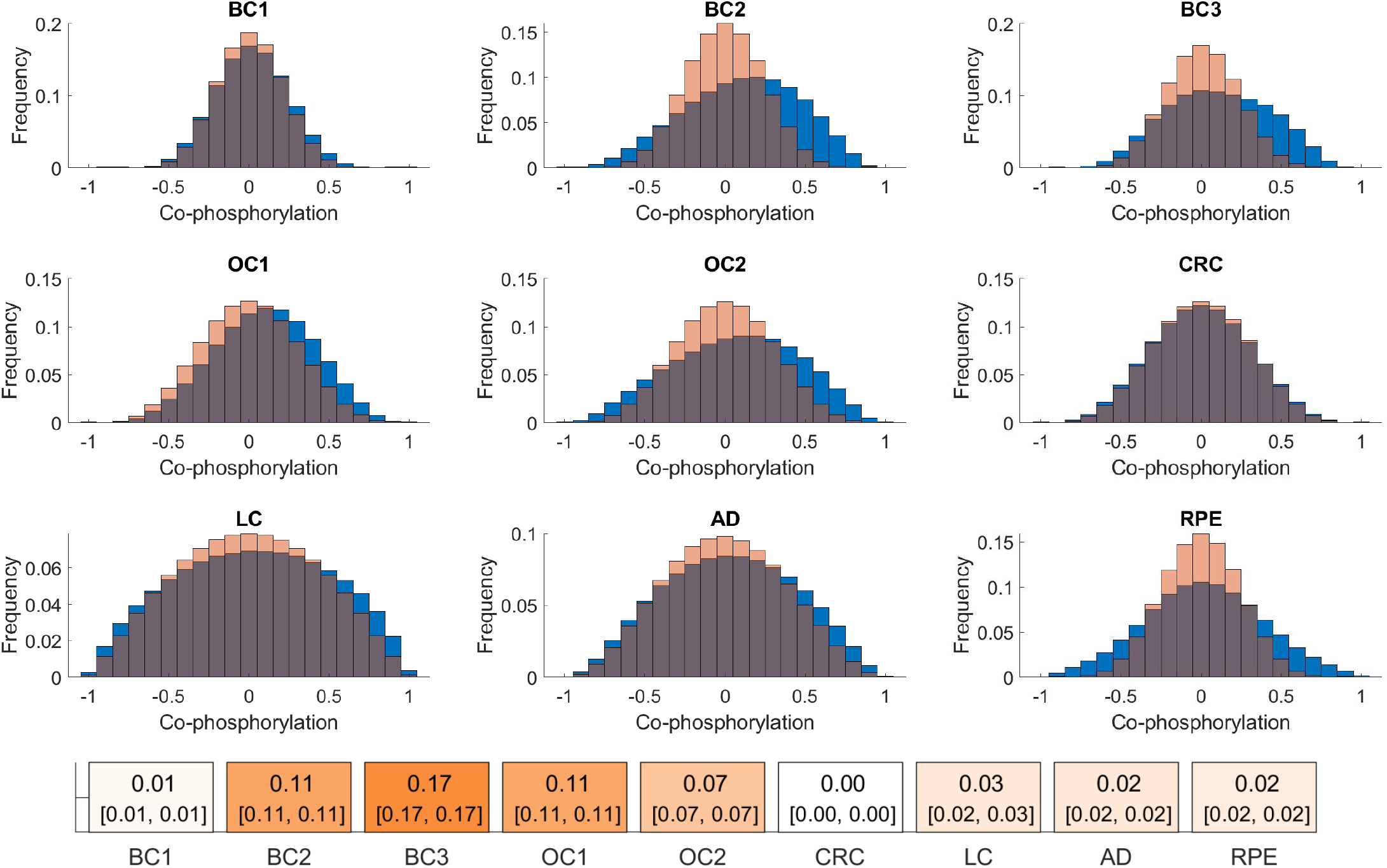
Statistical significance of co-phosphorylation. Each panel compares the distribution of co-phosphorylation computed on a specific dataset against that computed on randomly permuted data for each dataset. The blue histogram shows the distribution of co-phosphorylation (the correlation between the phosphorylation levels) of all pairs of phosphosites identified in the corresponding study, the pink histogram in each panel shows the average distribution of co-phosphorylation of all pairs of phosphosites across 100 permutation tests. The permutation tests are performed by randomly permuting all entries in the phosphorylation matrix. The difference between the means of each pair of distributions is given on the colored boxes below. The 95% confidence intervals for the difference are provided in brackets.

Furthermore, for most datasets (BC2, BC3, OC1), we observe that the mean co-P is clearly shifted to the right, as also indicated by the effect size and the significance of the t-statistic.. For other datasets (BC1, CRC), the difference between the means is close to zero and the corresponding t-statistics are less significant. However, even for these datasets, the KS-test indicates that the difference between the distributions is significantl, and visual inspection of the historgrams suggests that the histogram for observed Co-P values is always more spread. This observation suggests that these datasets also contain a large number of site pairs with negatively correlated phosphorylation levels. Clearly, as with positive correlation, negative correlation can also be indicative of a functional relationship between two phosphorylation sites

Taken together, for all studies considered, there are more pairs of phosphosites with (positively or negatively) correlated phosphorylation levels than would expected at random - hence a large fraction of these strong correlations likely stem from functional or structural relationships between the phosphosites.

### 3.2 Co-Phosphorylation of Intra-Protein Sites

Results of previous studies indicate that the phosphorylation of different sites of the same protein can lead to different functional outcomes [41, 42]. Here, with a view to characterizing the functional diversity of the phosphorylation sites on a single protein, we compare the Co-P distribution of pairs of phosphosites that reside on the same protein (intra-protein sites) against the Co-P distribution of pairs of phosphosites that reside on different proteins (inter-protein sites). We also investigate the effect of proximity between phosphorylation sites on the functional relationship between the sites. The results of this analysis are shown in Figure 2.

**Figure 2:**
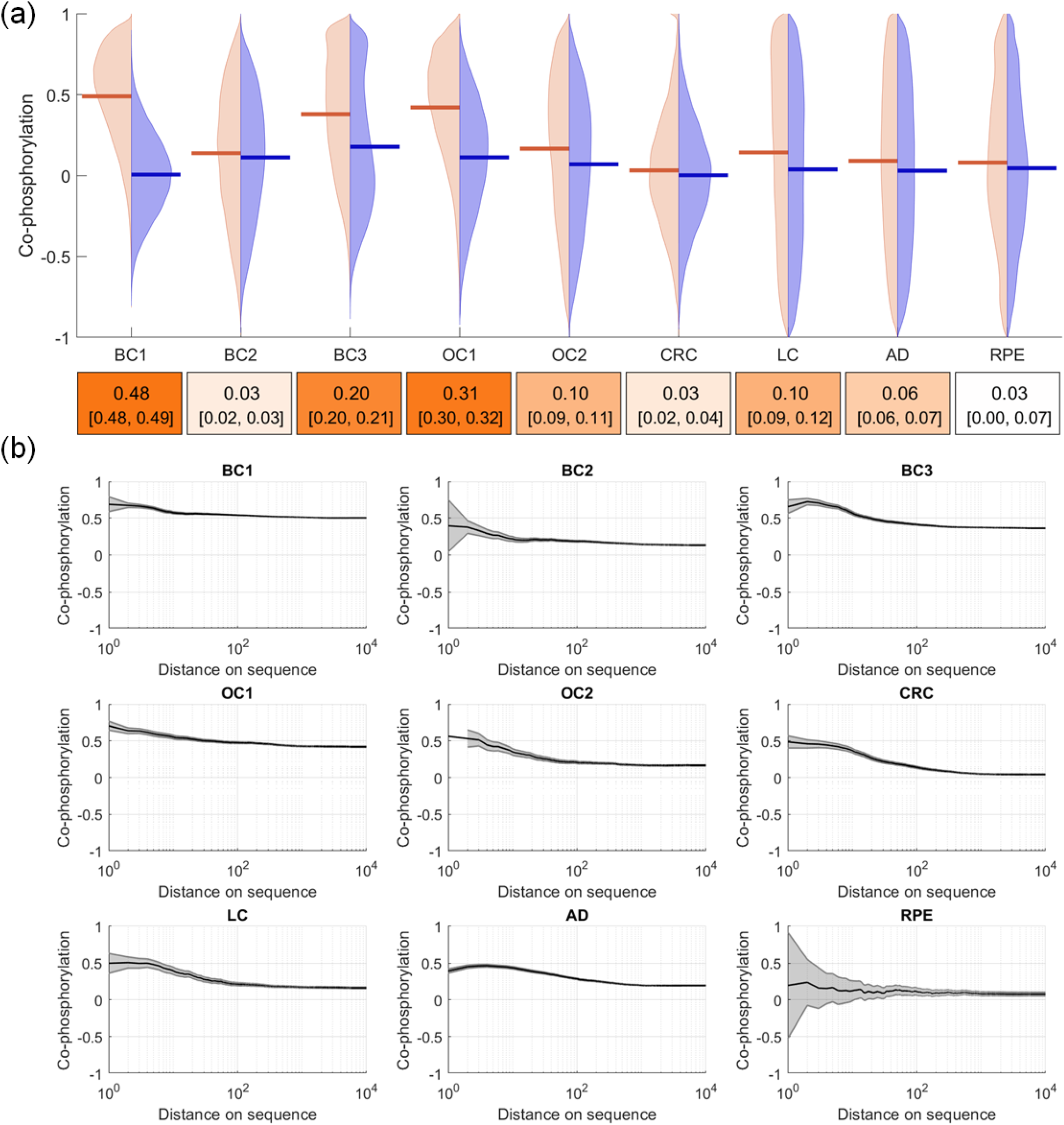
Co-phosphorylation of phosphorylation sites on the same protein. (a) Comparison of the distribution of Co-P for all site pairs that are on the same protein (orange histogram) vs. co-P for all pairs of sites on different proteins (blue histogram). Each violin plot represents a different dataset. Colored boxes below indicate the mean difference between the intra-protein pairs and inter-protein pairs. Within brackets, 95% confidence interval for the mean Co-P difference are provided. (b) The relationship between co-P and sequence proximity for pairs of sites that reside on the same protein. Each panel shows a different dataset, the x-axis in each panel shows the distance between sites on the protein sequence (in terms of number of residues) and the y-axis shows the co-phosphorylation between pairs of sites in close proximity (up to the corresponding distance in x-axis). The curve and shaded area respectively show the mean Co-P and its 95% confidence interval.

As seen in Figure 2(a), the distribution of co-Pforpairs of intra- and inter-protein sites are significantly different for most of the datasets (the mean differences and confidence intervals are provided in the figure, the p-values for the t-test as well as the KS-test are ≪ 1*E* −9 for all datasets except RPE). We consistently observe that the co-phosphorylation of intra-protein sites (orange histogram) is shifted towards high co-phosphorylation values. In other words,the phosphorylation levels of sites on the same protein are substantially more positively correlated as compared to the phosphorylation levels of sites on different proteins. While this observation can be partially explained by the impact of protein expression levels, a recent study showed that the protein abundance is overall not a strong indicator of phosphorylation fold-changes [43]. Thus, we hypothesize that intra-proteins pairs exhibit higher co-phosphorylation because those pairs are more likely to be targeted by the same kinase/phosphates, or that they are more likely to be functionally associated by being part of the same signaling pathways.

Note that, the differences between the datasets in terms of the difference of intra- and inter-protein pairs are highly prononunced (e.g., we observe strong difference for BC1, BC3, OC1 while difference is more modest for BC2, CRC, and AD). While there can be biological reasons for this difference, it is important to note that each of these datasets come from different platforms, different sample types (e.g., patient-derived xenografts vs. cell lines), different data collection procedures (e.g., protein degradation due to proteases in the sample), and are highly divergent interms of availability of data (number of identified sites and number of samples). For this reason, the observed differences between the datasets can also be attributed to experimental, technological, or statistical reasons. Further investigation is needed to elucidate potential biological differences between the systems that are represented by these datasets.

Next, we investigate whether the proximity on the protein sequence has any effect on the co-phosphorylation between two intra-protein sites. Since previous studies suggest that closely positioned sites tend to be phosphorylated by the same kinase [44], we expect a positive relation between sequence proximity and co-phosphorylation (i.e., we expect higher co-phosphorylation between close sites). To investigate this, we plot the relationship between the sequence proximity of intra-protein sites, and their co-phosphorylation (Co-P). Figure 2(b) shows that the closely positioned intra-protein sites have higher Co-P. Thus, we observe that as the phosphosites get far away from each other, their Co-P typically reduces.

### 3.3 Co-phosphorylation and Functional Association

Li et al. [23] show that phosphorylation sites that are modified together tend to participate in similar biological processes. Here, focusing on the dynamic range of phosphorylation, we hypothesize that phosphosite pairs with correlated phosphorylation profiles are likely to be functionally associated with each other. To test this hypothesis, we investigate the relationship between Co-P and a broad range of functional associations. Since our results in Figure 2 suggest that there is a considerable difference between intra-protein and inter-protein sites in terms of their co-phosphorylation, we perform stratified analyses for intra- and inter-protein pairs. The results of this analysis are shown in Figure 3.

**Figure 3:**
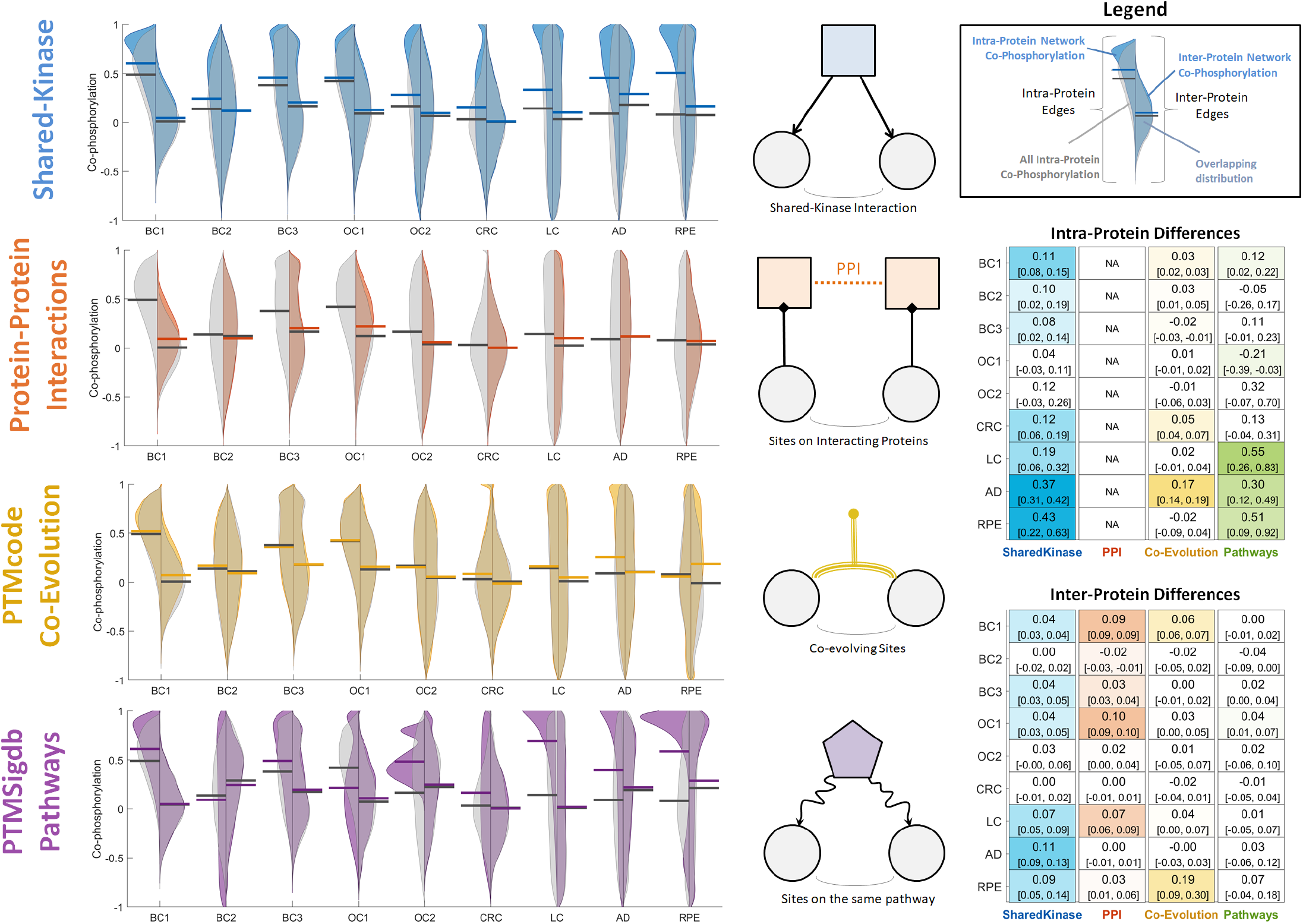
The relationship between co-phosphorylation and functional association between pairs of phosphorylation sites. In each panel, the violin plots compares the distribution of co-P for phosphosite pairs with an edge in the respective functional association network (colored histograms) against all phosphosite pairs (gray-colored histograms), across the 9 datasets that are considered. For each dataset, the left/right violin plots respectively show intra-/inter-protein pairs. The black horizontal lines show the mean Co-P for all (intra- or inter-protein) phosphosite pairs, the colored horizontal lines show the mean Co-P for functionally associated pairs. The four type of functional association networks that are considered are illustrated on the right side of the corresponding violin plot. On the rightmost side, the colored tables show the mean difference between functionally associated pairs and all phosphosite pairs (corresponding to the gap between colored and black horizontal lines in the violinplots) for 9 datasets and 4 functional networks. In each cell, the 95% confidence intervals for the mean difference is given within brackets.

#### Shared-Kinase Pairs

First, we consider the Co-P of the substrates of the same kinase (i.e., shared-kinase pairs) as annotated by PhosphositePlus. As seen in the Figure 3, in all datasets, the Co-P distribution of shared-kinase pairs is significantly shifted upwards, i.e., sites that are targeted by the same kinase are likely to exihibit stronger correlation of phosphorylation as compared to arbitrary pairs. While this difference is more pronounced for intra-protein pairs, it is also evident for inter-protein pairs. This observation is also in line with previous findings in the literature [21, 43].

#### Phosphorylation Sites on Interacting Proteins

Protein-Protein Interaction networks (PPI) encode physical and functional associations among proteins, thus have been used commonly for various inference tasks in cellular signaling. These tasks include identification of signaling pathways [45], identification of pathways that are mutated in cancers [46], prediction of the effect of mutations on protein interactions [47], and prediction of kinase-substrate associations [48]. It is also well-established that proteins that are coded by co-expressed genes are likely to interact with each other [49]. Here, we compare the PPI network and Co-P network to investigate the pattern of Co-P of pairs of phosphosites on interacting proteins. Note that, by definition, we only have this type of functional interaction for inter-protein sites. As seen in the Figure 3, in most of the datasets we consider (including BC1, BC3, OC1, OC2, LC, RPE), there is a clear upward shift of co-P for sites on interacting proteins. This suggests that sites on interacting proteins are likely to be co-phosphorylated. Identification of the specific protein-protein interactions (PPIs) that are associated with co-phosphorylation can be potentially useful in elucidating the mechanisms of these PPIs.

#### Co-evolution of Phosphorylation Sites

The conservation status of the phosphosites has been used as a tool to measure PTM activity [50]. It has been shown that co-evolving PTMs are likely to be functionally associated [51]. Here, we investigate the relationship between co-evolution and co-phosphorylation of phosphosites. The results of this analysis are shown in Figure 3. As seen in the figure, the association between co-evolution and co-phosphorylation is relatively weak compared to the association of co-P with other functional networks.

#### Phosphorylation Sites with Common Signaling Pathways

Identifying the signaling pathways that are dysregulated in any perturbation and disease is crucial for understanding the underlying mechanism of diseases. While most databases for signaling pathways are limited to gene or protein-centric information, PTMsigDB [35] provides a collection of PTM site-specific signatures that have been assembled and curated from public datasets. Using PTMsigDB, we investigate the Co-P of phosphosites that are involved in the same pathway. As seen in Figure 3, there is considerable difference between the Co-P distribution of the phosphosites that are involved in the same signaling pathway as compated to that of other phosphosite pairs. Similar to the results for shared-kinase pairs, this difference is more pronounced for intra-protein sites.

### 3.4 Predictive Power of Co-phosphorylation

Our results indicate that phosphosites involved in a common pathway or targeted by a common kinase are likely to be co-phosphorylated across different biological states. Motivated by this observation, we quantitatively assess the effectiveness of Co-P in predicting shared-kinase and shared-pathway associations between phosphorylation sites. While doing so, we also assess the contribution of Co-P evidence supported by multiple datasets to the reliability of predictions on functional association. For this purpose, we assess the predictive ability of Co-P computed using each individual dataset as well as the integrated Co-P computed using cross-dataset analysis. The results of this analysis are shown in Figure 4.

**Figure 4:**
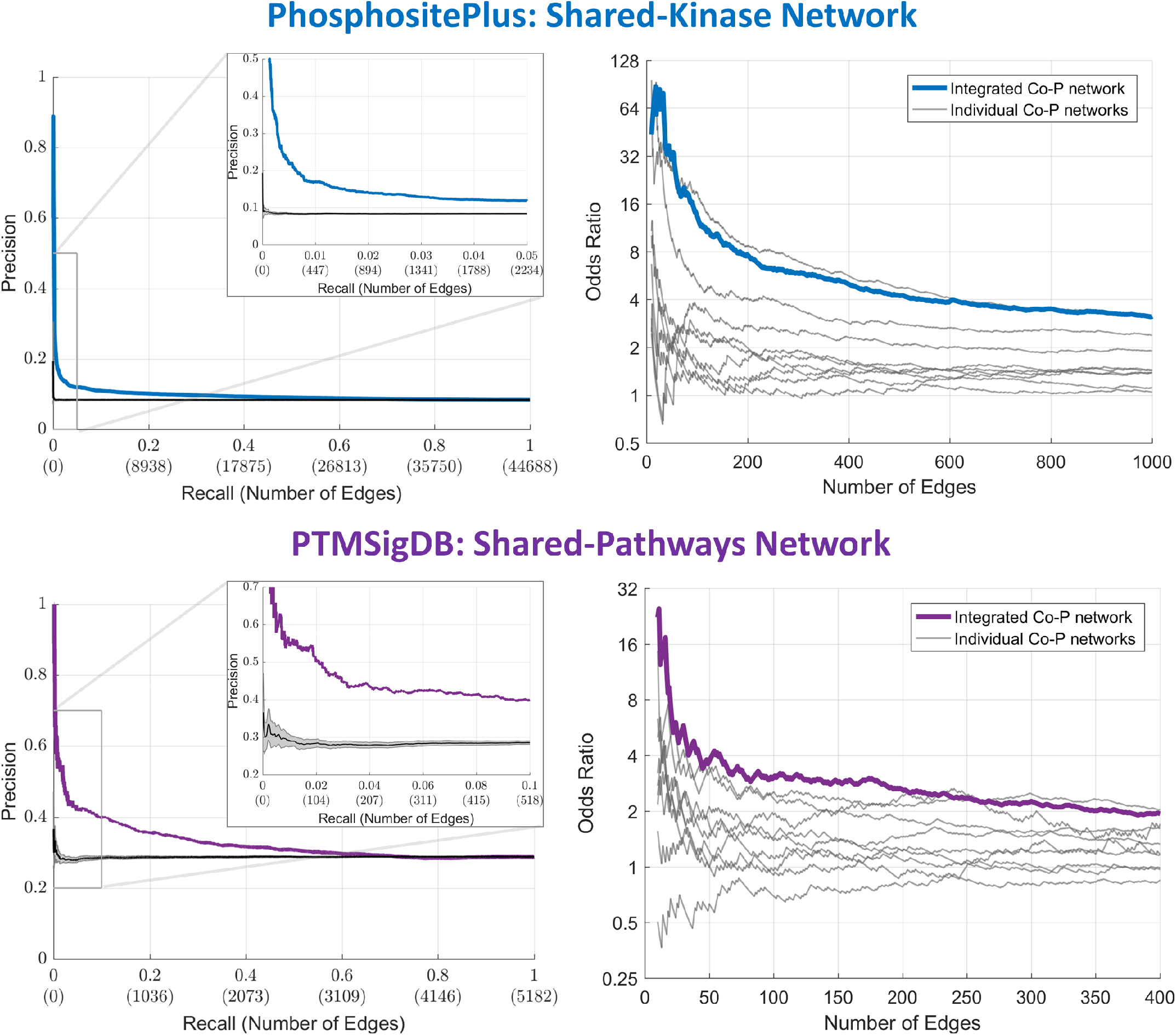
The utility of co-phosphorylation in predicting the functional association of phosphorylation sites. (Left) Precision-Recall curve showing the functional predictivity of the Co-P network obtained by integrating 9 different phospho-proteomic datasets. The shaded gray area shows the 95% confidence interval for the mean precision-recall curve for permutation tests obtained by randomly ranking pairs of phosphosites (across 20 runs). (Right) Comparison of the predictive performance of the integrated Co-P network against the 9 individual Co-P networks obtained using each dataset separately. The x-axis shows the number of pairs that are included in the co-P network, the y-axis shows the odds ratio of being connected in the respective functional network given that the sites are connected in the co-P network. (Top) Predicting shared-kinase associations. (Bottom) Predicting shared-pathway associations.

While constructing the co-P networks, we compute a co-P score for each pair of phosphosites, namely *c_d_*(*i, j*) for individual dataset *d* and *c_integrated_*(*i, j*) for the integrated network. We then sort the pairs according to this co-P score and apply a moving threshold to generate a series of co-P networks with increasing number of edges. In the left panel of Figure 4, the precision-recall curves for the ability of this network in predicting shared-kinase interactions (top-left panel) and shared-pathway interactions (bottom-left panel) are shown. In this context, recall is the defined as the fraction of edges in the corresponding functional network that also exist in the co-P network, whereas precision is defined as the fraction of edges in the co-P network that also exist in the functional network. To provide a baseline for the predictive ability of the co-P network, we also visualize the mean precision and 95% confidence interval for given recall for a random ranking of phosphosite pairs across 20 runs. As seen in the figure, the precision provided by the co-P network is significantly higher than random ordering for both functional networks. We also observe that co-P delivers higher precision for the shared-pathway network as compared to the shared-kinase network. This is likely because the information in PTMSigDB is sparser than the information in PhosphositePLUS.

The right panel of Figure 4 shows the odds ratio of a pair of sites being connected in the functional network as a function of the number of edges in the co-P network. Namely, in these plots, a point on the x axis corresponds to a co-P network with a given number of edges. For this network, the value on the y-axis shows the odds ratio of the event that two sites are connected in the functional network given that they are connected in the co-P network, as compared to a random pair of sites. As seen in the figure, for both shared-kinase and shared-pathway networks, the odds-ratio provided by the integrated co-P network is consistently higher than that provided by any individual network. While the odds-ratio of sharing a kinase goes up to 100 and the odds-ratio of being involved in the same pathway goes up to 30 for pairs of sites with co-P, these odds-ratios respectively converge to 4 and 2 as more edges are added to the integrated co-P network. Overall, these results suggest that co-P networks provide valuable information on the functional association of phosphorylation sites and this information becomes more reliable as co-P information from more datasets are included in the co-P network.

## 4 Conclusion

Mass-spectrometry techniques are advancing and more MS-based quantitative phosphoproteomics data are generated at high volumes. However, integration of these data may be challenging since the data is generated in different labs and in different contexts. By focusing on the relationships between pairs of phosphosites as opposed to their individual phosphorylation levels, co-phosphorylation networks can alleviate the dependency of computational and statistical methods on these factors. In this paper, we systematically investigated the relationship between co-phosphorylation and broad range of known functional associations between proteins and phosphorylation sites. Our results showed that the sites that are functionally associated tend to exhibit higher levels of co-phosphorylation. Our results also showed that the integration of co-phosphorylation networks across different datasets can improve the predictivity of co-phosphorylation, as compared to analyzing the datasets in isolation. These results highlight the power of network models and network-based analyses of phosphorylation data in predicting the functional relationships among phospho-proteins, kinases, and phosphatases in the context of cellular signaling.

## 5 Acknowledgement

This work was supported by National Institutes of Health grant R01-LM012980 from the National Library of Medicine. The content is solely the responsibility of the authors and does not necessarily represent the official views of the National Institutes of Health.

## References

[1] Mathias Wilhelm, Judith Schlegl, Hannes Hahne, Amin Moghaddas Gholami, Marcus Lieberenz, Mikhail M Savitski, Emanuel Ziegler, Lars Butzmann, Siegfried Gessulat, Harald Marx, et al. Mass-spectrometry-based draft of the human proteome. Nature, 509(7502):582–587, 2014.

[2] Klarisa Rikova, Ailan Guo, Qingfu Zeng, Anthony Possemato, Jian Yu, Herbert Haack, Julie Nardone, Kimberly Lee, Cynthia Reeves, Yu Li, et al. Global survey of phosphotyrosine signaling identifies oncogenic kinases in lung cancer. Cell, 131(6):1190–1203, 2007.

[3] Philip Cohen. The role of protein phosphorylation in human health and disease. the sir hans krebs medal lecture. European journal of biochemistry, 268(19):5001–5010, 2001.

[4] Vincentius A Halim, Monica Alvarez-Fernandez, Yan Juan Xu, Melinda Aprelia, Henk WP van den Toorn, Albert JR Heck, Shabaz Mohammed, and Rene H Medema. Comparative phosphoproteomic analysis of checkpoint recovery identifies new regulators of the dna damage response. Sci. Signal., 6(272):rs9–rs9, 2013.

[5] Sreenath V Sharma, Daphne W Bell, Jeffrey Settleman, and Daniel A Haber. Epidermal growth factor receptor mutations in lung cancer. Nature Reviews Cancer, 7(3):169–181, 2007.

[6] Doriano Fabbro, Sandra W Cowan-Jacob, Henrik Möbitz, and Georg Martiny-Baron. Targeting cancer with small-molecular-weight kinase inhibitors. In Kinase Inhibitors, pages 1–34. Springer, 2012.

[7] Peng Wu, Thomas E Nielsen, and Mads H Clausen. Small-molecule kinase inhibitors: an analysis of fda-approved drugs. Drug discovery today, 21(1):5–10, 2016.

[8] Serhan Yılmaz, Marzieh Ayati, Daniela Schlatzer, A Ercüment Çiçek, Mark R Chance, and Mehmet Koyutürk. Robust inference of kinase activity using functional networks. Nature communications, 12(1):1–12, 2021.

[9] Özgün Babur, Augustin Luna, Anil Korkut, Funda Durupinar, Metin Can Siper, Ugur Dogrusoz, Alvaro Sebastian Vaca Jacome, Ryan Peckner, Karen E. Christianson, Jacob D. Jaffe, Paul T. Spellman, Joseph E. Aslan, Chris Sander, and Emek Demir. Causal interactions from proteomic profiles: Molecular data meet pathway knowledge. Patterns, page 100257, 2021.

[10] James E Butrynski, David R D’Adamo, Jason L Hornick, Paola Dal Cin, Cristina R Antonescu, Suresh C Jhanwar, Marc Ladanyi, Marzia Capelletti, Scott J Rodig, Nikhil Ramaiya, et al. Crizotinib in alk-rearranged inflammatory myofibroblastic tumor. New England Journal of Medicine, 363(18):1727–1733, 2010.

[11] Caicun Zhou, Yi-Long Wu, Gongyan Chen, Jifeng Feng, Xiao-Qing Liu, Changli Wang, Shucai Zhang, Jie Wang, Songwen Zhou, Shengxiang Ren, et al. Erlotinib versus chemotherapy as first-line treatment for patients with advanced egfr mutation-positive non-small-cell lung cancer (optimal, ctong-0802): a multicentre, open-label, randomised, phase 3 study. The lancet oncology, 12(8):735–742, 2011.

[12] Joerg Neddens, Magdalena Temmel, Stefanie Flunkert, Bianca Kerschbaumer, Christina Hoeller, Tina Loeffler, Vera Niederkofler, Guenther Daum, Johannes Attems, and Birgit Hutter-Paier. Phosphorylation of different tau sites during progression of alzheimer’s disease. Acta neuropathologica communications, 6(1):52, 2018.

[13] Fumika Koyano, Kei Okatsu, Hidetaka Kosako, Yasushi Tamura, Etsu Go, Mayumi Kimura, Yoko Kimura, Hikaru Tsuchiya, Hidehito Yoshihara, Takatsugu Hirokawa, et al. Ubiquitin is phosphorylated by pink1 to activate parkin. Nature, 510(7503):162–166, 2014.

[14] Jang Hyun Choi, Alexander S Banks, Jennifer L Estall, Shingo Kajimura, Pontus Boström, Dina Laznik, Jorge L Ruas, Michael J Chalmers, Theodore M Kamenecka, Matthias Blüher, et al. Anti-diabetic drugs inhibit obesity-linked phosphorylation of pparγ by cdk5. Nature, 466(7305):451–456, 2010.

[15] Puneet Puri, Faridoddin Mirshahi, Onpan Cheung, Ramesh Natarajan, James W Maher, John M Kellum, and Arun J Sanyal. Activation and dysregulation of the unfolded protein response in nonalcoholic fatty liver disease. Gastroenterology, 134(2):568–576, 2008.

[16] Claudia Hernandez-Armenta, David Ochoa, Emanuel Gonçalves, Julio Saez-Rodriguez, and Pedro Beltrao. Bench-marking substrate-based kinase activity inference using phosphoproteomic data. Bioinformatics, 33(12):1845–1851, 2017.

[17] Noah Dephoure, Kathleen L Gould, Steven P Gygi, and Douglas R Kellogg. Mapping and analysis of phosphorylation sites: a quick guide for cell biologists. Molecular biology of the cell, 24(5):535–542, 2013.

[18] John R Yates III, Shabaz Mohammed, and Albert JR Heck. Phosphoproteomics, 2014.

[19] Yu Liu and Mark R Chance. Integrating phosphoproteomics in systems biology. Computational and structural biotechnology journal, 10(17):90–97, 2014.

[20] Elise J. Needham, Benjamin L. Parker, Timur Burykin, David E. James, and Sean J. Humphrey. Illuminating the dark phosphoproteome. Science Signaling, 12(565), 2019.

[21] Marzieh Ayati, Danica Wiredja, Daniela Schlatzer, Sean Maxwell, Ming Li, Mehmet Koyutürk, and Mark R Chance. Cophosk: A method for comprehensive kinase substrate annotation using co-phosphorylation analysis. PLoS computational biology, 15(2):e1006678, 2019.

[22] Marzieh Ayati, Mark R Chance,and Mehmet Koyuturk. Co-phosphorylation network sreveal subtype-specific signaling modules in breast cancer. Bioinformatics (Oxford, England), page btaa678, 2020.

[23] Ying Li, Xueya Zhou, Zichao Zhai, and Tingting Li. Co-occurring protein phosphorylation are functionally associated. PLoS computational biology, 13(5):e1005502, 2017.

[24] Kuan-lin Huang, Shunqiang Li, Philipp Mertins, Song Cao, Harsha P Gunawardena, Kelly V Ruggles, DR Mani, Karl R Clauser, Maki Tanioka, Jerry Usary, et al. Proteogenomic integration reveals therapeutic targets in breast cancer xenografts. Nature communications, 8(1):1–17, 2017.

[25] Philipp Mertins, Feng Yang, Tao Liu, DR Mani, Vladislav A Petyuk, Michael A Gillette, Karl R Clauser, Jana W Qiao, Marina A Gritsenko, Ronald J Moore, et al. Ischemia in tumors induces early and sustained phosphorylation changes in stress kinase pathways but does not affect global protein levels. Molecular & cellular proteomics, 13(7):1690–1704, 2014.

[26] Philipp Mertins, DR Mani, Kelly V Ruggles, Michael A Gillette, Karl R Clauser, Pei Wang, Xianlong Wang, Jana W Qiao, Song Cao, Francesca Petralia, et al. Proteogenomics connects somatic mutations to signalling in breast cancer. Nature, 534(7605):55–62, 2016.

[27] Hui Zhang, Tao Liu, Zhen Zhang, Samuel H Payne, Bai Zhang, Jason E McDermott, Jian-Ying Zhou, Vladislav A Petyuk, Li Chen, Debjit Ray, et al. Integrated proteogenomic characterization of human high-grade serous ovarian cancer. Cell, 166(3):755–765, 2016.

[28] Yuichi Abe, Maiko Nagano, Takahisa Kuga, Asa Tada, Junko Isoyama, Jun Adachi, and Takeshi Tomonaga. Deep phospho-and phosphotyrosine proteomics identified active kinases and phosphorylation networks in colorectal cancer cell lines resistant to cetuximab. Scientific reports, 7(1):1–12, 2017.

[29] Danica Wiredja. Phosphoproteomic Characterization of Systems-Wide Differential Signaling Induced by Small Molecule PP2A Activation. PhD thesis, Case Western Reserve University, 2018.

[30] Eric B Dammer, Andrew K Lee, Duc M Duong, Marla Gearing, James J Lah, Allan I Levey, and Nicholas T Seyfried. Quantitative phosphoproteomics of alzheimer’s disease reveals cross-talk between kinases and small heat shock proteins. Proteomics, 15(2-3):508–519, 2015.

[31] Cheng-Kang Chiang, Aleksander Tworak, Brian M Kevany, Bo Xu, Janice Mayne, Zhibin Ning, Daniel Figeys, and Krzysztof Palczewski. Quantitative phosphoproteomics reveals involvement of multiple signaling pathways in early phagocytosis by the retinal pigmented epithelium. Journal of Biological Chemistry, 292(48):19826–19839, 2017.

[32] Peter V Hornbeck, Bin Zhang, Beth Murray, Jon M Kornhauser, Vaughan Latham, and Elzbieta Skrzypek. Phosphositeplus, 2014: mutations, ptms and recalibrations. Nucleic acids research, 43(D1):D512–D520, 2015.

[33] Damian Szklarczyk, Andrea Franceschini, Stefan Wyder, Kristoffer Forslund, Davide Heller, Jaime Huerta-Cepas, Milan Simonovic, Alexander Roth, Alberto Santos, Kalliopi P Tsafou, et al. String v10: protein-protein interaction networks, integrated over the tree of life. Nucleic acids research, 43(D1):D447–D452, 2014.

[34] Pablo Minguez, Ivica Letunic, Luca Parca, Luz Garcia-Alonso, Joaquin Dopazo, Jaime Huerta-Cepas, and Peer Bork. Ptmcode v2: a resource for functional associations of post-translational modifications within and between proteins. Nucleic acids research, 43(D1):D494–D502, 2015.

[35] Karsten Krug, Philipp Mertins, Bin Zhang, Peter Hornbeck, Rajesh Raju, Rushdy Ahmad, Matthew Szucs, Filip Mundt, Dominique Forestier, Judit Jane-Valbuena, et al. A curated resource for phosphosite-specific signature analysis. Molecular & cellular proteomics, 18(3):576–593, 2019.

[36] Scott L Carter, Christian M Brechbühler, Michael Griffin, and Andrew T Bond. Gene co-expression network topology provides a framework for molecular characterization of cellular state. Bioinformatics, 20(14):2242–2250, 2004.

[37] Sara Ballouz, Wim Verleyen, and Jesse Gillis. Guidance for rna-seq co-expression network construction and analysis: safety in numbers. Bioinformatics, 31(13):2123–2130, 2015.

[38] Lin Song, Peter Langfelder, and Steve Horvath. Comparison of co-expression measures: mutual information, correlation, and model based indices. BMC bioinformatics, 13(1):328, 2012.

[39] Patrick E Meyer, Frederic Lafitte, and Gianluca Bontempi. minet: Ar/bioconductor package for inferring large transcriptional networks using mutual information. BMC bioinformatics, 9(1):461, 2008.

[40] Jeremy Miles. R squared, adjusted r squared. Wiley StatsRef: Statistics Reference Online, 2014.

[41] Hafumi Nishi, Alexey Shaytan, and Anna R Panchenko. Physicochemical mechanisms of protein regulation by phosphorylation. Frontiers in genetics, 5:270, 2014.

[42] Hafumi Nishi, Emek Demir, and Anna R Panchenko. Crosstalk between signaling pathways provided by single and multiple protein phosphorylation sites. Journal of molecular biology, 427(2):511–520, 2015.

[43] Osama A Arshad, Vincent Danna, Vladislav A Petyuk, Paul D Piehowski, Tao Liu, Karin D Rodland, and Jason E McDermott. An integrative analysis of tumor proteomic and phosphoproteomic profiles to examine the relationships between kinase activity and phosphorylation. Molecular & Cellular Proteomics, 18(8 suppl 1):S26–S36, 2019.

[44] Regev Schweiger and Michal Linial. Cooperativity within proximal phosphorylation sites is revealed from large-scale proteomics data. Biology direct, 5(1):6, 2010.

[45] Mitchell J Wagner, Aditya Pratapa, and TM Murali. Reconstructing signaling pathways using regular language constrained paths. Bioinformatics, 35(14):i624–i633, 2019.

[46] Matthew Ruffalo, Mehmet Koyutürk, and Roded Sharan. Network-based integration of disparate omic data to identify” silent players”in cancer. PLoS computational biology, 11(12):e1004595, 2015.

[47] Carlos HM Rodrigues, Yoochan Myung, Douglas EV Pires, and David B Ascher. mcsm-ppi2: predicting the effects of mutations on protein-protein interactions. Nucleic acids research, 47(W1):W338–W344, 2019.

[48] Heiko Horn, Erwin M Schoof, Jinho Kim, Xavier Robin, Martin L Miller, Francesca Diella, Anita Palma, Gianni Cesareni, Lars Juhl Jensen, and Rune Linding. Kinomexplorer: an integrated platform for kinome biology studies. Nature methods, 11(6):603–604, 2014.

[49] Arun K Ramani, Zhihua Li, G Traver Hart, Mark W Carlson, Daniel R Boutz, and Edward M Marcotte. A map of human protein interactions derived from co-expression of human mrnas and their orthologs. Molecular systems biology, 4(1):180, 2008.

[50] Jos Boekhorst, Bas van Breukelen, Albert JR Heck, and Berend Snel. Comparative phosphoproteomics reveals evolutionary and functional conservation of phosphorylation across eukaryotes. Genome biology, 9(10):R144, 2008.

[51] Pablo Minguez, Luca Parca, Francesca Diella, Daniel R Mende, Runjun Kumar, Manuela Helmer-Citterich, Anne-Claude Gavin, Vera Van Noort, and Peer Bork. Deciphering a global network of functionally associated post-translational modifications. Molecular systems biology, 8(1), 2012.

